# The Glucocorticoid Receptor is Required for Efficient Aldosterone-Induced Transcription by the Mineralocorticoid Receptor

**DOI:** 10.1101/2023.01.26.525745

**Authors:** Thomas A. Johnson, Gregory Fettweis, Kaustubh Wagh, Brian Almeida-Prieto, Manan Krishnamurthy, Arpita Upadhyaya, Gordon L. Hager, Diego Alvarez de la Rosa

## Abstract

The glucocorticoid and mineralocorticoid receptors (GR and MR, respectively) have distinct, yet overlapping physiological and pathophysiological functions. There are indications that both receptors interact functionally and physically, but the precise role of this interdependence is poorly understood. Here, we analyzed the impact of GR co-expression on MR genome-wide chromatin binding and transcriptional responses to aldosterone and glucocorticoids, both physiological ligands of this receptor. Our data show that GR co-expression alters MR genome-wide binding to consensus DNA sequences in a locus-and ligand-specific way. MR binding to consensus DNA sequences is affected by GR. Transcriptional responses of MR in the absence of GR are weak and show poor correlation with chromatin binding. In contrast, co-expression of GR potentiates MR-mediated transcription, particularly in response to aldosterone. Finally, single-molecule tracking of MR suggests that the presence of GR contributes to productive binding of MR/aldosterone complexes to chromatin. Together, our data indicate that co-expression of GR potentiates aldosterone-mediated MR transcriptional activity, even in the absence of glucocorticoids.

## INTRODUCTION

Adrenal glands coordinate physiological responses to cope with stress, acute injury or prolonged deprivation of water and food. Two important categories of adrenal hormones mediate key specific homeostatic responses: glucocorticoids (cortisol and corticosterone) and mineralocorticoids (aldosterone). However, these hormones show significant promiscuity. An excess of glucocorticoid signaling produces mineralocorticoid-like effects, particularly hypertension (1). Conversely, an excess of mineralocorticoids can mimic glucocorticoid effects, such as glucose homeostasis dysregulation and development of metabolic syndrome (2). The molecular basis for this cross-talk is at least partially explained by the close evolutionary relationship between the mineralocorticoid receptor (MR) and the glucocorticoid receptor (GR), which confers poor ligand specificity and overlapping modes of action (3, 4). Both mineralocorticoid and glucocorticoid hormones potently activate MR (5). Since glucocorticoids circulate at concentrations several orders of magnitude higher than aldosterone, certain cells co-express MR with 11-β-hydroxysteroid dehydrogenase type 2 (11-β-HSD2), an enzyme that metabolizes glucocorticoids into their biologically-inactive 11-keto metabolites, creating a low-glucocorticoid milieu (6). In contrast to MR, GR is partially selective, with potent activation by glucocorticoids and weak activation by mineralocorticoids (7). GR expression is essentially ubiquitous, while MR expression is more restricted and generally at lower abundance, except in the hippocampus and aldosterone-target epithelia such as the renal collecting duct and distal colon, where MR and GR abundance is similar (8). Pharmacological approaches or the use of mouse models with selective knockout of MR or GR in tissues that co-express both receptors conclusively demonstrate mutual MR and GR influence in determining glucocorticoid signaling outcomes (9-11). This, together with co-expression or not of 11-β-HSD2, generates at least three scenarios for corticosteroid hormone receptor function: GR-mediated responses to glucocorticoids; GR/MR-mediated responses to glucocorticoids; MR-mediated responses to aldosterone in the presence of presumably inactive GR.

Both MR and GR share a highly conserved DNA-binding domain (DBD), which implies that they recognize with high affinity the same DNA consensus sequence, known as “Hormone Response Element” (HRE) (12), and likely regulate a partially overlapping set of genes. The largest differences in the amino acid sequences of MR and GR proteins occur in the N-terminal domain (NTD) with *Mus musculus* MR containing over 100 more amino acids than *M. musculus* GR and only 29% similarity (Fig. S1A). By contrast, the DBD and the ligand-binding domain (LBD) of MR and GR have 90% and 69% similarity, respectively. The NTD of MR is important for gene regulation (13, 14). The AF1 activation regions of the MR NTD appear to be separated into two distinct amino acid sequences, AF1a and AF1b, similar to the androgen or progesterone receptors but unlike GR, which has one central AF1 domain in its NTD (14-16). In addition, MR is reported to have a region with intrinsic inhibitory function placed between AF1a and AF1b (16). The divergent structure of MR and GR NTDs may account to some extent for differential, tissue-specific transcriptional responses (17, 18).

To further complicate the picture, it has been conclusively demonstrated that MR and GR can physically interact to form heteromers (19-25). Examination of the functional properties of MR/GR interaction has produced conflicting experimental results. Gene-reporter assays or studying the expression of specific genes indicate that GR may enhance MR transcriptional activity in certain cell lines (26, 27), although it appears to be inhibitory or non-influential in others (19-21, 28). A recent study performed in keratinocytes demonstrated that MR co-expression alters GR genomic binding and modulates the global transcriptional response to the synthetic glucocorticoid dexamethasone (9). The fact that GR expression is ubiquitous and generally higher than MR (8) implies that MR will typically function in the presence of significant levels of GR. The functional effects of this co-expression are unclear. Data obtained *in vivo* suggest that both receptors may be needed for potent aldosterone biological effects (29, 30). However, there are no studies to date directly analyzing the global influence of GR on MR-mediated transcriptional responses, whether driven by aldosterone or glucocorticoids.

Given the physical interaction between both factors, the molecular basis for their specific physiological roles, overlapping functions and pathological consequences of dysregulation can only be understood after clearly defining the consequences of co-expression in genome-wide studies. To this end, we took advantage of a well-characterized cellular system to study MR chromatin binding, gene regulation, and single-molecule dynamics in the presence or absence of GR. Our results indicate that GR profoundly affects MR genome-wide chromatin binding in a locus-and ligand-specific way and generally potentiates MR/aldosterone-mediated gene transcription, correlating with apparent stabilization of MR productive chromatin binding.

## RESULTS

To study genome-wide function of mammalian MR both in the presence and absence of the closely related GR, we stably introduced a GFP-tagged MR into a well-characterized GR knockout (GRKO) C127 mouse cell line and its parental line that expresses GR endogenously (31). The GFP-tag of the integrated MR is inserted after residue 147 of the N-terminus of the receptor, which has been shown to optimize its hormone response over an N-terminal GFP tag and it readily translocates to the nucleus with hormone (Fig. S1B-C; (32)). We collected genome-wide data sets for RNA expression and chromatin immunoprecipitation (ChIP) from the two MR cell lines before and after hormone treatments with 10nM aldosterone (Aldo) or with 100nM corticosterone (Cort), both saturating conditions for MR (7). Additionally, the same chromatin preparations from the MR parental cells treated with Aldo or Cort were also used to ChIP endogenous GR. We also compared the MR hormone-mediated response to previously reported genome-wide wild type and mutant GR datasets from the same cell lines (31, 33).

### Co-expression of GR alters MR genome-wide binding in locus-specific and ligand-specific ways

We performed ChIP-Seq for MR in the GRKO cells and the parental cells to determine how the presence of GR may affect MR binding across the genome. We treated cells with vehicle, Aldo or Cort for 1 hour prior to sample collection to detect differences in MR chromatin binding with Aldo (MR specific) or with Cort, which activates both MR and GR (5, 34, 35). GRKO cells treated with Aldo have slightly stronger MR chromatin binding than with Cort, as seen by normalized ChIP-seq signal intensity (Fig. 1A, aggregate plots). A union list of 2403 called MR peaks in the GRKO cells distributes into three clusters: 164 Cort-specific peaks (cluster 1), 782 Aldo-specific peaks (cluster 2), and 1457 peaks that occur with either treatment (cluster 3). MR binding in the parental cells with endogenous GR stands in contrast to its binding in the GRKO cells, as Cort induces the strongest MR binding instead of Aldo (Fig. 1B, aggregate plots). A Union list of 1718 MR peaks in the parental cells distributes into two large clusters: 1213 Cort-specific peaks (cluster 4), 501 Aldo/Cort-shared peaks (cluster 5). Importantly, only a small group of only four (weak) Aldo-specific peaks were found (not shown).

**Figure 1.**
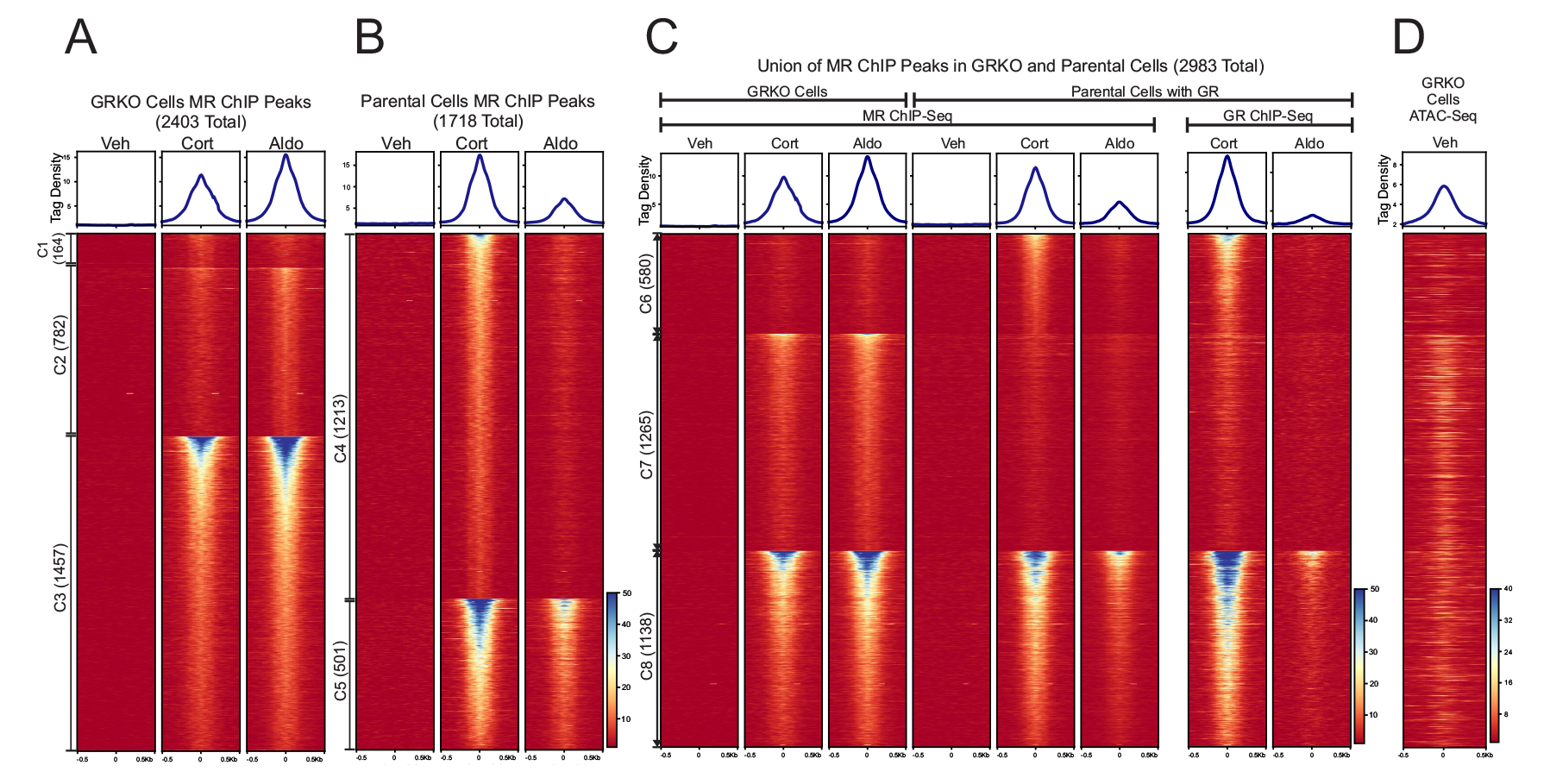
Chromatin binding of MR. **A-B**. Comparison of MR binding after 1 Hr. treatment with vehicle, 100nM Cort or 10nM Aldo in two cell lines (GRKO with no GR or Parental with endogenous GR). Aggregate plots represent total ChIP-Seq tag density of all peaks normalized as reads per genomic content (1x normalization). Heatmaps represents ± 500bp around the center of the MR peak. ChIP-Seq intensity scale is noted lower right on a linear scale. Clusters of peaks (C1-C5) are labeled on the left with the peak number in parentheses and are sorted from high to low signal for the condition with the highest overall signal. A small cluster of four Aldo-specific peaks in the parental cells are not shown on the heatmap. **C-D**. A union list of MR ChIP peaks from A-B was created (see methods) and clustered by cell line and hormone treatment. 2983 unique MR peaks are distributed into Parental-specific/Cort (C6), GRKO-specific/Aldo (C7) or shared between to two cell types (C8). Aggregate plots and heatmaps are displayed as described for A-B for both MR ChIP and endogenous GR ChIP (parental cells only). ATAC-seq data are from untreated GRKO cells with stably expressed GFP-GRwt (31). The ATAC data heatmap is sorted the same as ChIP data in C and intensity scale is noted lower right on a linear scale.

A comparison of genome-wide MR binding between the two cell lines indicates how much GR influences MR chromatin binding and how this influence is likely related to MR-GR receptor interactions. A union list of MR binding peaks across the 2 cell lines with either ligand has a total of 2983 unique peaks in 3 clusters (Fig. 1C, clusters 6-8; Table S1). The 2403 MR peaks in the GRKO cells distribute into clusters 7 and 8 while the 1718 MR peaks in the parental cells distribute into clusters 6 and 8. The Cort-treated parental cells have 580 MR peaks (cluster 6) that do not occur in the GRKO cells with either ligand, suggesting Cort-liganded GR is required to enable MR binding at these sites. This requirement for Cort-liganded GR does not necessarily signify direct interaction or interdependence of the two nuclear receptors. GR binding could simply cause chromatin accessibility changes at these sites which could then enable MR binding; however, binding site clusters 7 and 8 provide more compelling evidence for receptor interaction. Cluster 7 has 1265 peaks specific to the GRKO cell line, indicating that in the Parental cells Cort-or Aldo-liganded GR inhibits MR binding at these sites. The cluster 7 MR binding in the parental cells is even weaker with Aldo treatment, again anticorrelated with MR binding in the GRKO cells. The cluster 7 endogenous GR ChIP-Seq signal in the parental cells also has very low binding with Cort treatment and virtually no binding with Aldo, suggesting that Aldo-liganded GR cannot efficiently bind these sites. It is, thus, unlikely that GR is simply out-competing MR for these sites (36). The 1138 MR binding sites shared between the two cell lines (cluster 8) exhibit the strongest signal intensity; however, the sites show the weakest peak intensities when both receptors are present and liganded by Aldo. This observation suggests that after 10nM Aldo treatment, GR not only binds chromatin poorly but also inhibits MR binding. The lower binding of Aldo-liganded GR (versus Cort-liganded GR) is directly shown by ChIP-Seq of the parental cells with a GR antibody (Fig. 1D).

Taken together, these data suggest that liganded GR has a dominant effect on the binding of MR, both potentiating at some sites inaccessible to MR by itself (cluster 6) and reducing its binding at sites it can bind when acting in the absence of GR (clusters 7 and 8). These effects on MR binding at clusters 7 and 8 could be due to GR not completely translocating to the nucleus with Aldo, GR reducing chromatin accessibility of the binding site or, more likely, due to mixed receptor heteromers have different binding efficacies. Very few GR binding events have been shown to reduce chromatin accessibility at the site of direct receptor binding (37). It appears that the type of ligand bound to the interacting receptors, Aldo or Cort, also affects both receptors’ binding efficiencies, with Aldo reducing the binding efficiency of both MR and GR when the receptors occupy the nucleus together.

### Motif analyses show that MR binding favors consensus NR3C1-4/AP1 motifs and is affected by GR

We performed separate motif analyses on clusters 6-8 from the 2983 MR binding sites that occur across the two cell lines. We used various known motifs in the Homer database (38) that reflect the 13mer NR3C1-4 steroid receptor consensus sequence GnACAnnnTGTnC as a proxy for an MR binding motif (Table S2). The three Homer MR motifs range in the stringency to the above consensus sequence primarily at the positions 3 and 5 “A”, position 9 “T” or position 13 “C” while the other positions are well conserved.

The 580 cluster 6 MR peaks that only appear in the parental cells with Cort treatment had some form of MR-like binding motif between 37-62% of sites (Fig.2; Table S2). The prevalence of the consensus steroid hormone receptor motif likely enabled both GR and, subsequently, MR binding (36). The AP1-like nTGAnTCAn motif (Table S2; Fig. 2) occurred between 2-4% of sites while THRb, Runx1, ZNF domain and ETS motifs were also detected. The 1265 cluster 7 GRKO-specific MR peaks were less enriched for MR-like consensus motifs (11-26% of sites) while AP1- like motifs occurred more often at 8-15% of sites (Fig. 2; Table S2). The 1138 MR peaks of cluster 8 that are shared across the two cell lines were as enriched as cluster 6 for MR-like binding motifs (36-62% of sites) and similarly or slightly less enriched as cluster 7 for AP1-like motifs (5-10% of sites).

**Figure 2.**
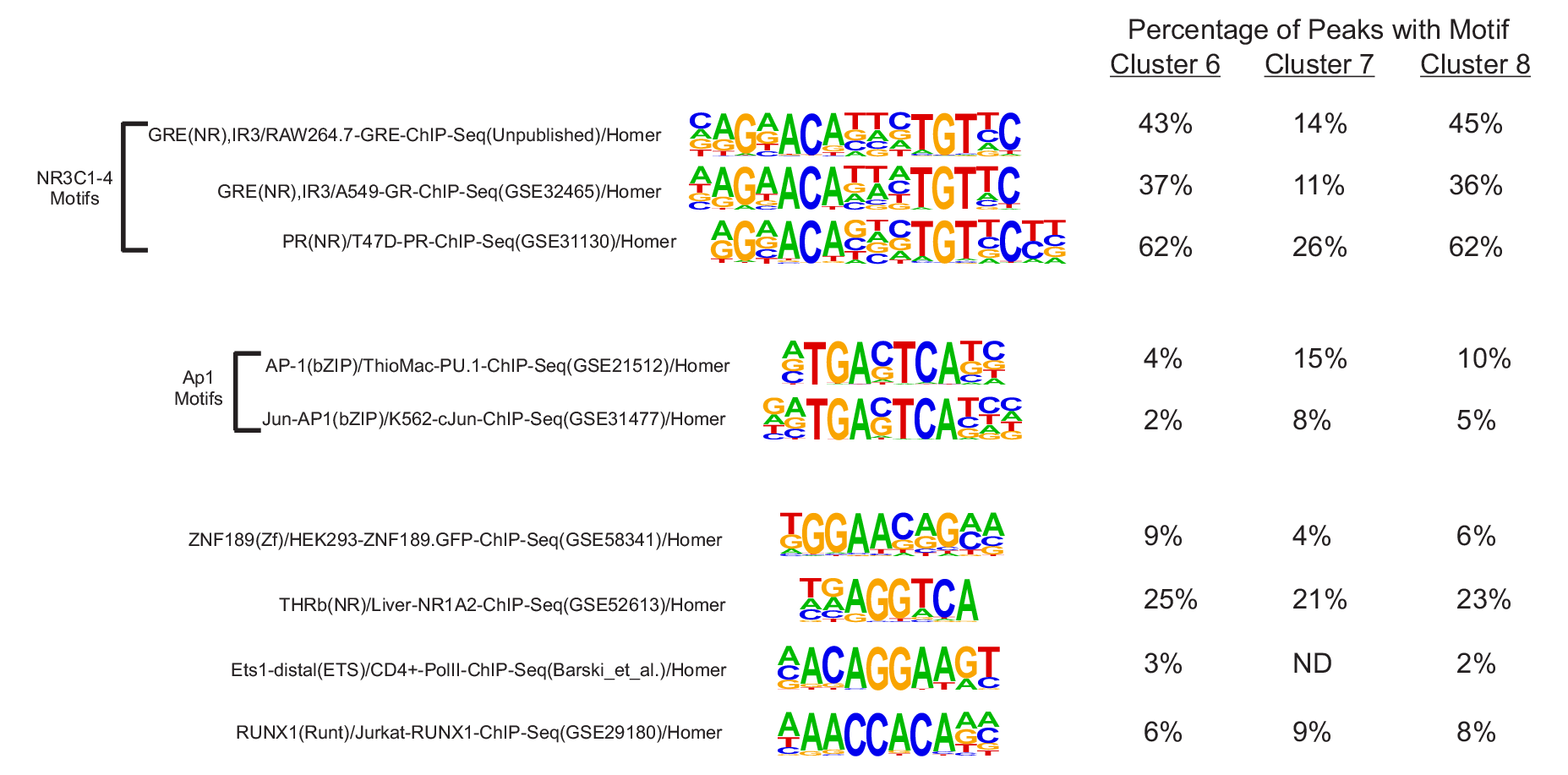
De novo motif analysis of MR binding. Figure shows position weight matrix logos for known motifs from the HOMER database. For Clusters 6-8, the percentage of MR binding sites that contain the designated known motif. See also Table S2 for numbers of sites containing a designated motif and background information.

The motif analyses align with the MR ChIP-seq data in that the GR-dependent cluster 6 MR peaks are more enriched than cluster 7 peaks for NR3C1-4 motifs. Our previous studies have shown GR to be capable of binding to inaccessible, nucleosomal sites prior to hormone with GRE consensus motifs (33, 39) whereas GRKO-specific sites (cluster 7) may more often have factors like AP1 bound prior to hormone that enable MR binding (40). This is reflected in previously published chromatin accessibility data of GRKO cells that show higher ATAC signal in cluster 7 prior to hormone than in cluster 6 (Fig. 1D; (31)). Cluster 8 MR sites exhibit the highest receptor binding intensity and have relatively higher enrichment of NR3C1-4 motifs than cluster 7 and higher AP1 consensus motifs than cluster 6. Like cluster 7, cluster 8 also exhibits higher pre-hormone accessibility than cluster 6 (Fig. 1D).

Overall, the motif analyses shows that the shared peaks (Cluster 8) contain both a higher frequency of MR-like motifs and more accessible chromatin prior to hormone. These characteristics likely explain why cluster 8 has the highest ChIP signal intensities. When MR is present alone in the GRKO cells, it binds more promiscuously at sites with fewer MR-like motifs (Cluster 7).

### MR exhibits weaker transcriptional activity without GR

To explore how receptor binding is related to changes in gene expression, we performed genome-wide total RNA sequencing (RNA-Seq) before and after hormone treatments in the GRKO and parental cell lines. We compared the differential expression (DE) analysis of vehicle versus 2 hours of exposure to either hormone from each cell line. We chose a false discovery rate (FDR) cutoff of 0.01 as determined by DESeq2 (via Homer) (41) to determine which genes show a change in exon RNA levels after treatment using two to three biological replicates per condition.

The parental cells with endogenous GR treated with Cort exhibit the strongest hormone response in both the number of up-or down-regulated genes and quantitative changes in RNA levels after treatment (Table S3). We categorized protein-coding genes that met our FDR cutoff and exhibited a hormone-dependent change in RNA levels of plus or minus Log2 0.5 or greater. The parental cells treated with Cort had 210 hormone-responsive genes compared to 53 genes when treated with Aldo, 52 of which are common to both hormone treatments (Fig. 3A). The GRKO cells exhibit a much-reduced response to both Cort and Aldo with 12 and 17 genes changed, respectively, 11 of which are common to both treatments and also overlap with the common responsive genes in the parental cells. Even among 30 genes shared between the two cell lines that meet the FDR criteria regardless of fold change, the hormone response in the GRKO cells is attenuated compared to the cells with GR (Fig. 3B). Among these 30 genes, the hormone dependent fold change in the GR expressing cell line is lower with Aldo treatment compared to Cort; however, in the GRKO cells the level of hormone response is similarly weak for either hormone compared to the gene responses in the parental cells. This indicates MR by itself is a poor transcriptional regulator with either Cort or Aldo. The higher gene response of the parental cells with Cort treatment can be primarily attributed to GR, its natural receptor. The number of Cort-responsive genes is similar to that obtained in these cells when treated with dexamethasone (33). Most importantly, the gene response of the parental cells is also greater for Aldo treatment suggesting that MR regulates genes better with its natural ligand in concert with Aldo-liganded GR. RNA-seq data from Aldo-treated parental cells without MR show only 4 genes that meet the FDR and FC cutoffs, none of which overlap with hormone-responsive genes in the MR expressing cell lines. (Fig. 3A; Table S3). This shows that GR by itself cannot induce a significant transcriptional response when liganded to Aldo, only in conjunction with MR.

**Figure 3.**
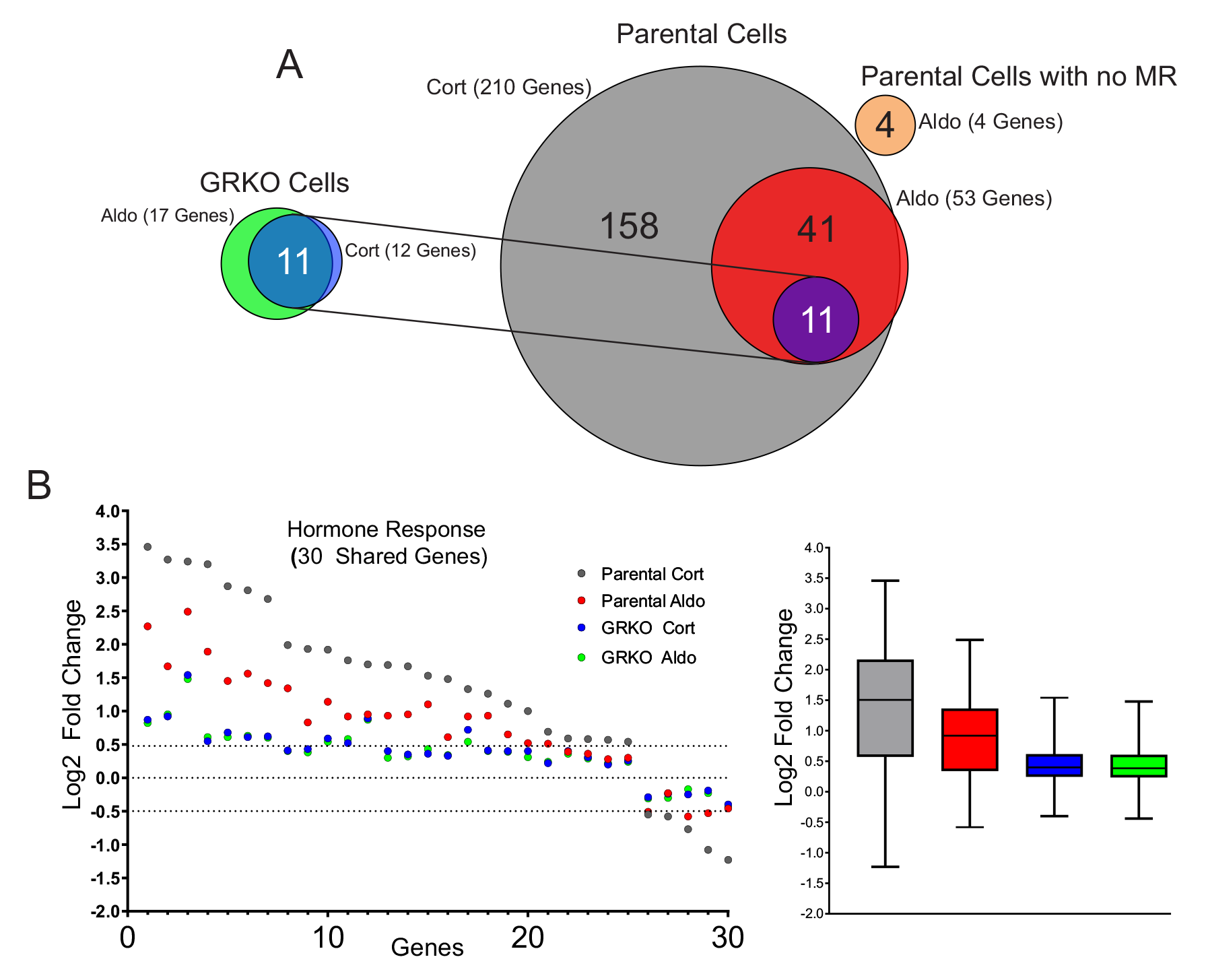
MR hormone response in the presence or absence of GR. **A.** Venn diagrams of hormone-regulated protein-coding genes (2 Hrs. treatment/vehicle) to 100nM Cort or 10nM Aldo. Total number of hormone responsive genes (FDR ≤ 0.01, Log2 FC ≥ +/- 0.5) denoted in parentheses for MR expressing GRKO cells, MR expressing Parental cells or Parental cells without MR. Circles connected with lines denote 11 common hormone-responsive genes to the two cell lines. **B.** Scatter plot of Log2 FC for 30 genes common (meet FDR 0.01 cutoff only) to the four denoted conditions. Box and whiskers plot of the same data display interquartile range (IQR) depicting the 25th, 50th and 75th percentile as box with the median as black bar. The whiskers mark the most induced and repressed genes.

### Intergenic eRNAs correlate with MR-mediated gene transcription

The RNA-Seq data are not in agreement with the MR and GR ChIP-Seq results discussed above if we consider overall ChIP signal (binding intensity and number of peaks) to correlate with gene response (42). Among the four experimental conditions (two cell lines and two hormones), the parental line treated with Aldo generates the fewest MR and GR peaks but exhibited the second strongest transcriptional response. The GRKO cell line generates a comparatively weak transcriptional response to both Aldo and Cort despite producing more ChIP peaks than the respective treatments in the GR-containing parental line. Within each cell line, the ChIP and the RNA results do correlate as Aldo induces slightly more genes and more ChIP peaks in the GRKO cells while Cort does the same in the parental line.

We used Homer to annotate the MR peaks in the three clusters of Fig. 1C to the closest gene *in cis* and detected if these closest genes are among the 211 MR-responsive genes of the parental cells. Among the 1138 GRKO/Parental shared peaks of cluster 8, 79 were closest to hormone responsive genes, while 28 and 32 peaks were closest to hormone responsive genes in Clusters 6 and 7, respectively (Table S1). These results are in line with those obtained by Ueda et al., which identified 25 out of 1414 ChIP-seq peaks placed proximal to aldosterone-regulated genes, as determined by microarray analysis, including common MR/GR targets such as SGK1, GILZ or Tns1 (43). Hormone responsive genes were sometimes detected near peaks from more than one cluster (ex. Tns1, Glul) or even all three clusters (ex. Ampd3, Tgm2). Of the 11 common MR responsive genes in GRKO cells, 10 are linked to annotated peaks in clusters 1, 2 or 3. The remaining gene, CTGF, a known MR/Aldo target gene in the heart (44), has a nearby distal peak in cluster 3, but with a non-hormone responsive gene occurring closer to it detected by the Homer peak annotation. MR binding at several loci near a hormone responsive gene likely contributes to its transcriptional response and these loci may occur in more than one of the clusters in Fig. 1B. However, the strongest MR binding sites (cluster 8), as measured by ChIP signal intensity, are associated in *cis* to the most hormone responsive genes.

Active enhancers produce short bidirectional RNAs, known as enhancer RNAs (eRNAs), at sites of transcription factor binding and actively transcribing genes (45, 46). We used the total RNA-Seq data to look for eRNAs at intergenic MR ChIP peaks as an indication of such activity. We left out MR peaks annotated to intron, UTR, exon and promoter sites to avoid RNA signals made by transcription near or within gene bodies. Of the 2983 MR peaks that make up clusters 6 to 8, 1576 are classified as intergenic according to Homer. We plotted the intergenic eRNA signal at these peaks sorted as a subset of each cluster in Fig. 1C and by ChIP signal intensity (Fig. 4). The eRNA heatmaps show little hormone-dependent change in signal at intergenic peaks in the GRKO cells; while increases in signal can be observed in the parental cells, more so with Cort than Aldo. In the parental cells, the increase in eRNA levels correlate with MR ChIP signal intensity at all three clusters, with Cluster 8.1 showing more change in overall eRNA signal than either cluster 6.1 or cluster 7.1. Thus, the eRNA signal correlates with MR ChIP binding signal in the Parental cells with GR present, but not in the GRKO cells. The overall eRNA signal also correlates with the overall transcriptional response at the gene level.

**Figure 4.**
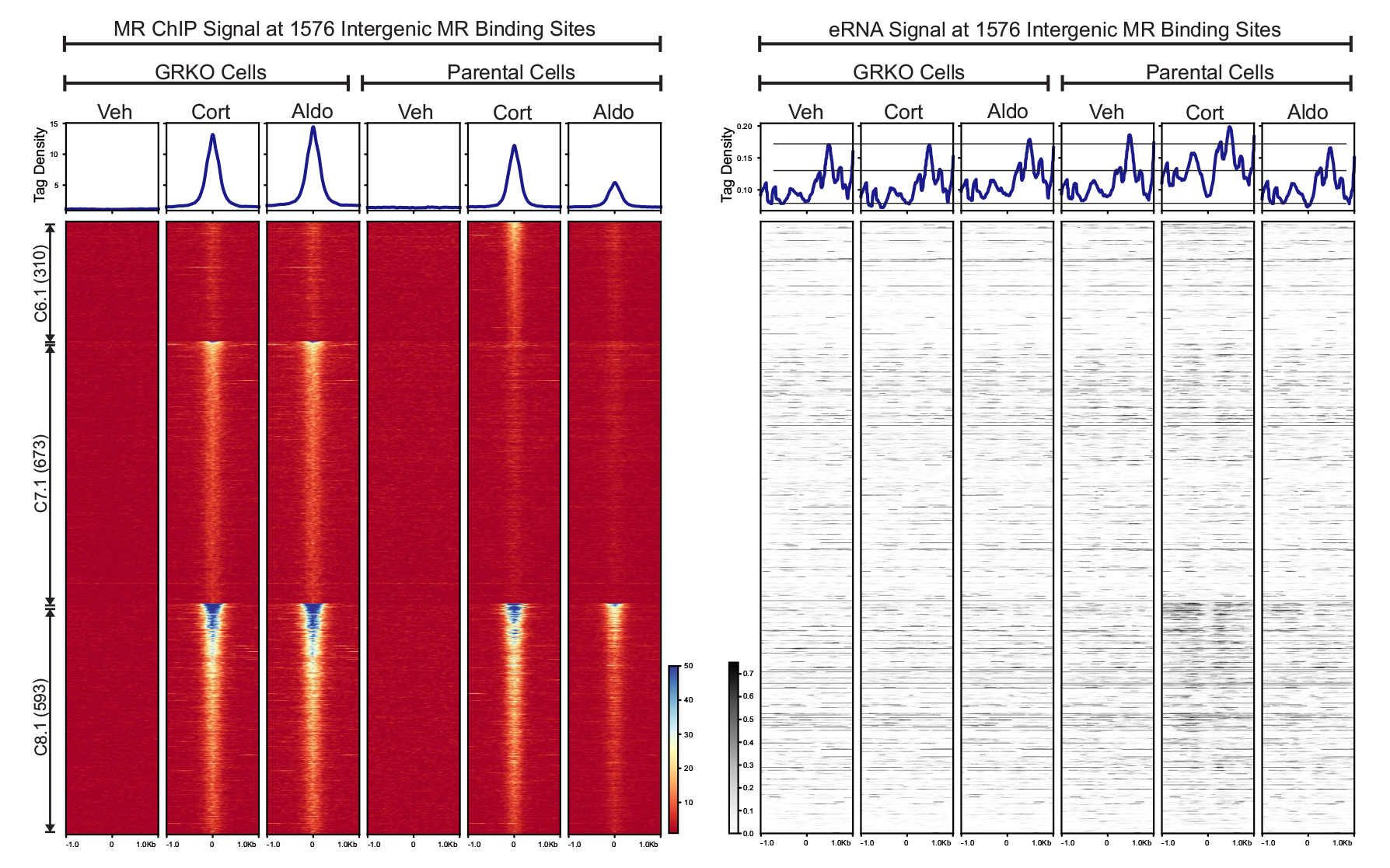
Enhancer RNA signals at intergenic MR chromatin binding peaks. The left heatmap shows subsets of clusters 6-8 (Fig 1C) representing intergenic MR ChIP peaks. Aggregate plots represent total ChIP-Seq tag density of all peaks normalized as reads per genomic content (1x normalization). Heatmaps represents ± 1kb around the center of the MR peak. ChIP-Seq intensity scale is noted lower right on a linear scale. Clusters of peaks are labeled on the left with the peak number in parentheses and are sorted from high to low signal for the condition with the highest overall signal. The right heatmap shows total normalized RNA-seq signal from merged replicates in the same order and breadth as the MR-ChIP heatmaps. Aggregate plots represent total RNA-Seq tag density normalized as reads per genomic content (1x normalization). RNA-Seq intensity scale is noted lower left on a linear scale.

### The MR NTD contributes to transcriptional activity

The transcriptomic data demonstrate that MR binding alone (ChIP signal intensity) in the absence of GR is not sufficient to induce a robust transcriptional response at the gene level or even of eRNAs at sites of MR binding. This could be due to a few possibilities. (*i*) MR requires a cofactor that may be present in cell types that naturally express MR but is missing in our cell lines. However, the extent of the MR gene response to Aldo in the parental cells argues against this being a wide-ranging limitation to MR activity. (*ii*) MR requires the involvement of GR and works better in concert with it to produce a transcriptional response. The greater hormone response of the parental cells that express endogenous GR agrees with this inference. (*iii*) The unique structure of the NTD of MR may convey an inhibitory effect on steroid responsive genes similar to that shown by Litwack and colleagues on the MR-responsive Na/K ATPase β1 gene (28) or through its recruitment of particular co-repressors as shown by Lombes and colleagues (16).

To further explore how the NTD of MR affects its genome-wide hormone response, we performed RNA-Seq using an NTD-truncation mutant (MR-580C) in both the GRKO and Parental cell lines (Fig. 5A). We categorized protein-coding genes that met our FDR cutoff (0.01) and exhibited a hormone-dependent change in RNA levels of plus or minus Log2 0.5 or greater FC. Like the full-length version of MR (MRwt), the MR-580C mutant exhibited a weak gene response in the GRKO cells with 11 and 20 MR-responsive genes to Cort and Aldo treatment, respectively (Fig. 5B; Table S3). The presence of endogenous GR in the Parental cells appears to potentiate the transcriptional effects of the MR-580C. The Cort-treated Parental cells have 314 responsive genes that meet the FDR and FC cutoffs while the Aldo-treated have 41 genes that meet the cutoffs. Again, the larger number of responsive genes with Cort versus Aldo can be attributed mainly to the presence of endogenous GR, but MR-580C activates/represses more genes with Aldo treatment in the presence of GR than in the GRKO cells (Fig. 5B; Table S3). A comparison of 218 genes Cort-responsive genes meeting only the FDR cutoff and common to the MRwt cells and MR-580C cells show overall similar hormone responses (Fig. 5C). A similar comparison of 59 genes with Aldo treatment often shows reduced gene responses with MR-580C compared to MRwt (Fig. 5D). These data suggest that the NTD of MR is indeed functional in our model cell lines and contributing to the hormone-dependent transcriptional response, especially with Aldo treatment in the presence of GR. The MR-580C data also indicate that the NTD of MR does not have an overall inhibitory effect on hormone-dependent transcriptional activity, as MRwt response is as high or higher than MR-580C.

**Figure 5.**
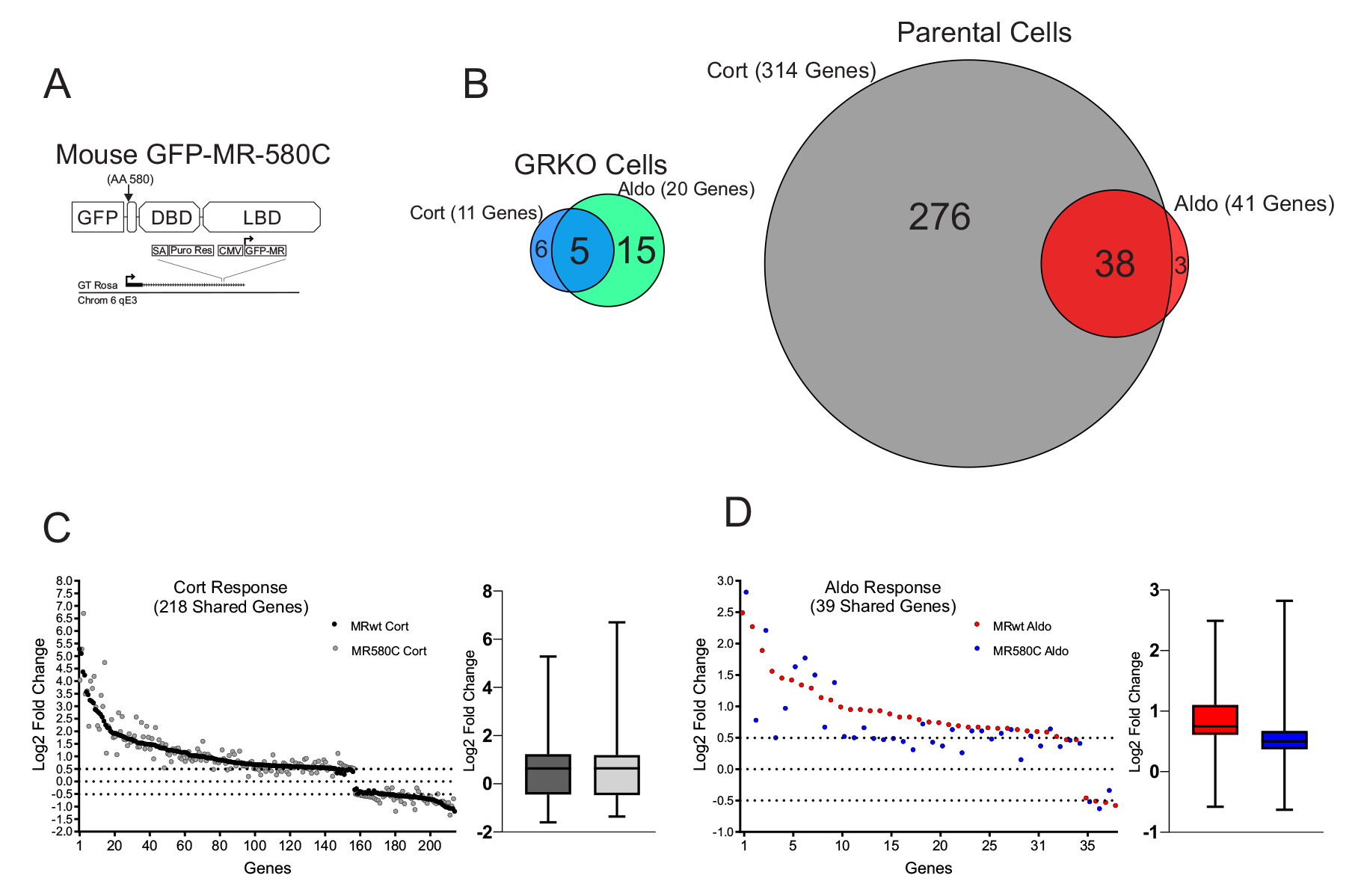
Contribution of MR N-terminal domain to the transcriptional response. **A.** Gene diagram of the stably expressed the MR NTD mutant. **B.** Venn diagrams of hormone-regulated protein-coding genes (2 Hrs. treatment/vehicle) to 100nM Cort or 10nM Aldo. Total number of hormone responsive genes (FDR ≤ 0.01, Log2 FC ≥ +/- 0.5) denoted in parentheses for MR expressing GRKO cells or MR expressing Parental cells (with endogenous GR). Circles connected with lines denote common hormone-responsive genes to the two cell lines. **C.** Scatter plot of Log2 FC for 218 Cort-responsive genes in the Parental cells common (meet FDR 0.01 cutoff only) to MRwt (Fig.2A) and MR580C. Box and whiskers plot of the same data display interquartile range (IQR) depicting the 25th, 50th and 75th percentile as box with the median as black bar. The whiskers mark the most induced and repressed genes. **D.** Scatter and Box and Whiskers plots of Log2 FC for 39 Aldo-responsive genes in the Parental cells common to MRwt and MR580C, as described above.

### Single molecule tracking of MR suggests GR contributes to productive binding of MR-Aldo complexes

Given that our overall ChIP-seq results do not explain well the global changes in MR-mediated transcription induced by GR co-expression, we tested whether GR-induced changes in MR chromatin binding and transcriptional activity correlate with altered receptor dynamics in the nucleus. We performed single-molecule tracking (SMT) of transiently transfected HaloTag-MR chimeras in the GRKO and Parental cells to determine the spatiotemporal dynamics of MR under these conditions. We fluorescently labeled Halo-MR with low concentrations of organic dye (see methods; (47)), and imaged cell nuclei using highly inclined laminated optical sheet (HILO) microscopy (48). We focused on the spatial mobility of molecules that remain stable on the order of tens of seconds as these have been shown to be correlated with transcriptional outcomes (49). We imaged the cells every 200 milliseconds (to minimize photobleaching), with 10 millisecond exposures (to minimize motion blur) (50). We note that at this frame rate, freely diffusing molecules will rapidly exit the focal plane on our analysis timescales. This method allows us to track molecules that are bound to chromatin. The temporal projection of representative SMT movies along with overlaid tracks are shown in Fig. 6A.

**Figure 6.**
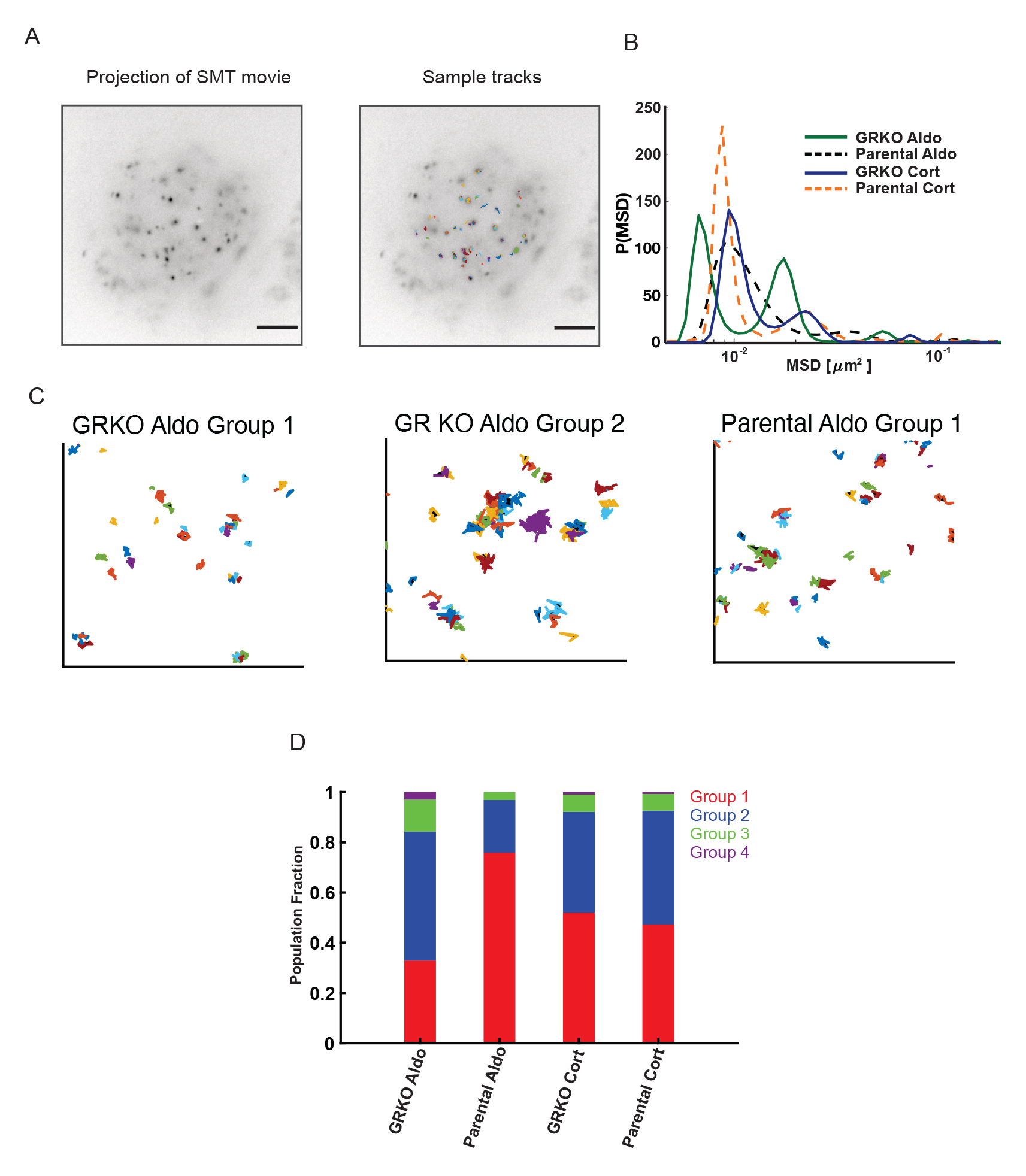
Single molecule tracking of MR. **A.** Temporal projection of an MR single molecule tracking movie (left) overlaid with tracks (right). Scale bar 5 μm **B.** Distribution of MR mean squared displacement (MSD) at a timelag of 0.8 s, obtained by iteratively fitting the van Hove correlation (vHc) function using the Richardson-Lucy algorithm for each of the four conditions – GRKO-Aldo (green solid line), Parental Aldo (black dashed line), GRKO-Cort (blue solid line), Parental-Cort (orange dashed line). **C.** Representative tracks for MR in the dominant lowest mobility groups. (Left) MR-Aldo group 1 tracks in GRKO cells, (center) MR-Aldo group 2 tracks in GRKO cells, (right) MR-Aldo group 1 tracks in Parental cells. Note that MR-Aldo in Parental cells exhibits a very small fraction of group 2 tracks (see also panel D). **D.** Population fractions for the different mobility groups for the indicated condition. N_cells_/N_tracks_: 51/1878 (GRKO MR-Aldo), 51/2000 (GRKO MR-Cort), 57/2419 (Parental MR-Aldo), 60/2036 (Parental MR-Cort). Two biological replicates were collected for each condition.

Recent SMT studies have identified two distinct mobility groups for chromatin and chromatin-bound transcriptional regulators (51-53). Transcriptionally active steroid receptors and other transcription factors show a substantially higher proportion of binding in the lowest mobility state and binding in this state requires an intact DNA-binding domain as well as domains necessary to recruit cofactors (53). We hypothesized that the substantial gene response to Aldo in Parental cells could result from an increased association of MR in the lowest mobility group. To test this, we collected SMT data of MR under the two different hormone stimulation conditions in both GRKO and Parental cell lines. We then iteratively fit the jump distance histogram at a specified time lag using an algorithm developed by Richardson (54) and Lucy (55) to uncover the distribution of mean-squared displacements (MSD) or equivalently, the distribution of diffusivities (see Methods). Applying this analysis to MR trajectories for the four different conditions, we find that single MR molecules exhibit multiple mobility groups exemplified by the distinct peaks in the MSD distribution (Fig. 6B). We can use the MSD distribution to classify the ensemble of MR trajectories into multiple mobility groups (Fig S2; see Methods). As can be seen from the areas under the peaks, most MR tracks distribute into the two lowest mobility groups (Fig, 6B). Representative tracks for the two lowest mobility groups are shown for MR-Aldo in the GRKO cells (Fig. 6C, left and center) and MR-Aldo in the Parental cells (Fig. 6C, right). Strikingly, MR-Aldo in the Parental cells exhibits a substantially higher proportion of molecules in the lowest mobility group as compared to any other condition (Fig. 6D). While the proportion of MR-Cort in the lowest mobility group is higher than MR-Aldo in the GRKO cell line, the presence of GR in the Parental cell line does not affect these proportions. Taken together, the presence of GR leads to an ∼2.5 fold increase in the population fraction of MR-Aldo in the lowest mobility group as compared to that in the absence of GR (Fig. 6D). Binding in this lowest mobility group is markedly higher for other steroid receptors (GR, androgen, progesterone, and estrogen receptors) in their transcriptionally active (liganded) state as compared to that in their inactive state (53). Even though MR-Aldo presents the least binding as measured by ChIP-seq (Fig. 1A, B), our live cell studies suggest that a higher proportion of MR binding in the transcriptionally active group (group 1) could contribute to the higher transcriptional output of Aldo-treated Parental cells as compared to Aldo-and Cort-treated GRKO cells (Fig. 2). However, we cannot discount the presence of additional mechanisms such as differential co-regulator recruitment.

## DISCUSSION

In this study, we have demonstrated the global importance of GR to enhance aldosterone-driven transcriptional function of MR. Because both receptors respond to Cort, we cannot distinguish the contribution of MR/GR interactions with this treatment using RNA-seq alone. We have shown that when liganded to Aldo or Cort, MR by itself binds to hormone response elements but cannot efficiently elicit a transcriptional response. In contrast, when acting in the presence of GR, Aldo-liganded MR can increase transcription of genes and at intergenic enhancers that it cannot efficiently induce on its own. Further, GR by itself cannot elicit a significant gene response when liganded to Aldo, where the hormone acts much like an antagonist by binding receptor without imparting a functional response (56). Thus, in the presence of Aldo, both receptors appear to act together, enhancing MR-mediated gene response. This correlates with GR-dependent altered single molecule dynamics of MR, with Aldo-liganded MR showing an ∼2.5-fold increase in the population fraction of group 1 in the presence of GR (Fig. 6D).

The MR/GR coactivation of genes occurs despite lower overall levels of genomic binding by both receptors when present in the nucleus together compared to MR alone, specifically with Aldo treatment. Aldo-induced MR ChIP-seq peak intensities are reduced in all 3 clusters in the presence of GR (parental cells) versus its absence (GRKO cells), as shown in Fig. 1B. This suggests that genomic binding of MR (as measured by ChIP) is not sufficient to promote a full gene response. GR also has much reduced binding when liganded to Aldo suggesting that the two receptors are not competing for binding at the same response elements, in agreement with previous results obtained using forebrain-specific MR knockout mice (57). When Cort is the ligand, the level of MR binding increases simultaneously with GR binding and is higher than binding with Aldo. This may indicate that interaction of the two sister-receptors likely imposes its effects by the recruitment of co-regulators necessary for modulating transcription and not via higher levels of binding. Having a hetero-multimer of Aldo-liganded GR and MR may more effectively recruit co-factors than MR can accomplish by itself. Further studies on co-factor recruitment and transcriptional response are needed to answer this question.

Starting with the original studies by Trapp et al. (25) and Liu et al. (21), it has long been known that MR and GR are able to form heterocomplexes, although the functional impact of this interaction has been elusive (19-25). The functional effect of GR on MR action has been mainly studied in the context of glucocorticoid signaling, based on the common assumption that in the presence of 11-β-HSD2, local glucocorticoid levels are very low and thus GR would be inactive and not affect Aldo-mediated MR activity. Reporter gene transactivation assays using low levels of cortisol stimulation (up to 10 nM, thus favoring MR over GR binding) showed increased transcriptional responses when both receptors were present (25). However, this result seems to be dependent on promoter context, since opposite results were obtained with a reporter assay using a different promoter (21). Evidence for a direct MR/GR interaction was later expanded to a negative GRE (nGRE), with data suggesting that heteromerization of MR and GR directly mediates corticosteroid-induced trans-repression of the 5-HT1A receptor promoter (26). Further work performed with rainbow trout MR and GR receptors using gene reporter assays suggested that MR-GR interaction may be involved in cortisol responses, with a dominant-negative role of MR in the process (58). Interestingly, this study found that MR inhibitory role persists even in the presence of its antagonist eplerenone, suggesting that MR transcriptional activity is not important in the process. Other reports agree with this notion, where MR plays a dominant-negative role on GR-mediated glucocorticoid-regulated gene expression, further suggesting that the NTD of MR is the domain involved in this effect through heterodimerization (59). The importance of this domain is confirmed by our data, as shown in Fig. 5. Mifsud et al. tested the relevance of MR and GR interaction in a more physiological context, testing MR and GR binding to GREs of common glucocorticoid-target genes (Fkbp5, Per1 and Sgk1) in hippocampal neurons after exposure of rats to environmental stressors (60). Their results are consistent with gene-dependent binding of MR and GR to GREs as homo-and/or heterodimers. GR binding seemed to facilitate MR binding to GREs in Fkbp5 and Per1 genes under high-glucocorticoid conditions. Taken together, these studies generally indicate that MR and GR co-expression may impact glucocorticoid-mediated gene expression, but are limited by the lack of genome-wide binding or transcriptional analyses. More recently, Rivers et al investigated the global effect of MR on GR genomic binding in transfected neuroblastoma N2a cells using ChIP-nexus (24). Their results show that MR and GR bind to overlapping, highly similar sites (58% of them with GRE motifs). RT-qPCR experiments measuring expression of selected genes (Syt2, Sgk1, Dusp4 and Ddc) showed that MR expression alone produced modest changes in expression upon 100 nM Cort stimulation, while GR co-expression induced more potent changes. This last experiment does not allow differentiating between MR-mediated and GR-mediated transcriptional changes. The authors proposed a tethering mechanism where GR mediates MR interaction with chromatin (24).

Few studies have directly investigated the impact of GR on Aldo-mediated MR transcriptional activity. Tsugita et al. examined this question with neuroblastoma and colon carcinoma cell lines expressing MR in the absence or presence of co-transfected GR and using reporter gene assays (27). This study demonstrated a lack of Aldo-induced luciferase activity unless GR is co-transfected. This MR-rescuing effect is specific for GR, since other steroid receptors such as PR, or AR did not have any effect. Interestingly, mutations in the DBD of GR prevented the potentiation of MR activity, suggesting that GR DNA binding is critical for the effect (27).

What is the molecular basis for the modulation of MR transcriptional activity by GR? Our data indicates that global, steady-state binding of MR to chromatin is not predictive of transcriptional activity. Interestingly, MR seems to be intrinsically more stable in its interaction with DNA than GR, as shown by hormone washout experiments (23, 24), but this does not explain the changes in transcription seen upon GR co-expression. It may be argued that the proposed tethering mechanism, where GR mediates MR indirect binding to DNA may play a role in explaining our results (24). In this scenario, the MR ChIP peaks detected in our experiments in GRKO and parental cell lines would not be directly comparable, since the latter would correspond to a different mode of interaction that is more productive transcriptionally. However, the fact that GR binds DNA poorly when Aldo is the ligand but still has a prominent effect on potentiating MR activity rules out this possibility. Another intriguing possibility is that GR, by interacting with MR and modulating its final oligomeric state as we recently reported (61), changes MR conformation to a more active state. Interestingly, our SMT data supports the idea of GR-induced differences in the kinetics of MR interaction with DNA. This is consistent with a previous report showing higher *in vitro* stability of MR/GR-DNA complexes when compared to MR alone (25). The situation may be more complicated, since MR and GR appear to interact with a specific GRE (a known binding site in the Per1 gene) in a cyclical way, possibly alternating homo-and heterocomplexes (62). Unfortunately, this study did not address MR dynamic interaction with chromatin in cells where GR is absent, precluding a more detailed analysis of the impact of GR on MR kinetics. On the other hand, MR and GR’s cyclical interaction with chromatin also apply when Aldo is used as agonist (62), consistent with our findings at a genome-wide level. This further reinforces the idea that GR participates in modulating MR-mediated transcriptional responses even when Aldo is the agonist.

Our data suggest that MR has likely evolved to work in concert with its more transcriptionally active sister receptor, GR, which is present in most tissues, including those where MR plays important cellular functions. By studying Aldo-activated MR function in the presence and absence of GR we mimic MR function in mammalian tissues where GR is present but, via Cort inactivation by 11-β-HSD2, can bind only, or mainly to Aldo, which is likely a physiologically relevant scenario. Ackermann et al. showed that while MR is constitutively nuclear in the Aldo-sensitive distal nephron, GR responds to fluctuations in Aldo circulating levels, at least in rats (63). Specifically, when Aldo levels are lowered by dietary NaCl loading, GR is localized to the cytosol, while MR remains nuclear. It is necessary to abrogate Aldo synthesis by adrenalectomy to achieve cytosolic localization for both MR and GR (63). Given the high circulating glucocorticoid levels during the peak of the circadian rhythm, it is possible that small amounts of glucocorticoids reach MR, which has high affinity for them. However, low doses of glucocorticoids would not activate GR and therefore the situation would result in relatively low MR activity. Only an increase in Aldo, which would be sensed by MR and also partially by GR would result in a more prominent MR-mediated response. This is consistent with a mechanism where GR plays an important role in the Aldo response, as originally proposed by Geering et al. (30) and indirectly corroborated by experiments using targeted knockout of the MR in the renal collecting duct (64) or overexpression of the GR in the renal collecting duct (65). In general, progressive recruitment of GR may contribute to the modulation of MR in the Aldo-sensitive distal nephron. This mechanism may have an impact in situations of altered glucocorticoid and mineralocorticoid signaling, including those induced under pathological situations or by pharmacological treatment of patients.

It appears that, in addition to MR/GR, other combinations of receptors within the estrogen and steroid receptor subfamily (NR3) give rise to functionally altered heteromers. These “atypical” interactions, including association of GR with the progesterone, estrogen or androgen receptors (66, 67) are increasingly recognized as important factors in determining transcriptional outcomes of hormone signaling, although the underlying mechanisms remain poorly understood (66). The apparent stabilization of productive chromatin binding of MR by GR suggests a more general mechanism that may underly cross-talk between members of the NR3 subfamily of nuclear receptors.

## MATERIALS AND METHODS

### Plasmids constructs and mutagenesis

A fully functional mouse MR fluorescent derivative with insertion of eGFP after amino acid 147 has been previously described (32). eGFP-MR was subcloned in plasmid Donor-Rosa26_Puro_CMV (31), with CMV promoter-driven expression, a puromycin resistance cassette and homology recombination arms specific for the mouse Gt(ROSA)26Sor locus. pX330 CRISPR/Cas9 plasmid, containing a guide RNA sequence to target the Gt(ROSA)26Sor locus, was a gift from Feng Zhang (Addgene plasmid #42230; (68)). Halo-tagged MR was constructed using In-Fusion cloning. The entire NTD of MR was deleted using the Quickchange XL mutagenesis kit, generating construct MR-580C. All constructs and mutations were confirmed by DNA sequencing.

### Cell culture and generation of cell lines by CRISPR/cas9

Cell lines were grown in Dulbecco’s modified Eagle’s medium (DMEM, Gibco) supplemented with 5 μg/ml tetracycline (Sigma-Aldrich #T7660), 10% fetal bovine serum (Gemini), sodium pyruvate, nonessential amino acids, and 2 mM glutamine. Cells were maintained in a humidifier at 37C and 5% CO2. Cells were plated for experiments in DMEM supplemented with 10% charcoal/dextran-treated serum for 24 hrs prior to hormone treatment. Cell lines used in this study derive from mouse mammary carcinoma cell line C127 (RRID: CVCL_6550) cells. Knockout of endogenously-expressed GR generating GRKO cells has been previously described (31). Transient transfections were performed using Jetprime (Polyplus) according to the manufactureŕs instructions. eGFP-tagged MR was stably integrated in the genome using CRISPR/Cas9. To that end cells were co-transfected with pX330 CRISPR/Cas9 plasmid with a donor plasmid containing eGFP-MR driven by the CMV promoter. Donor plasmid insertion was selected by puromycin treatment followed by fluorescence-activated cell sorting (FACS). Expression of MR in sorted polyclonal cell declined with time and therefore we selected stable lines by one additional round of FACS, followed by single-cell cloning. GFP-MR expression in individual clones was confirmed by confocal microscopy and western blot using monoclonal antibody rMR1-18 1D5 (developed by Gomez-Sanchez et al. (69), and obtained from the Developmental Studies Hybridoma Bank, created by the NICHD, National Institutes of Health and maintained at The University of Iowa, Department of Biology) as previously described (20). MR agonists aldosterone and corticosterone were obtained from Sigma and dissolved in ethanol. Cells were plated for experiments in DMEM growth medium supplemented with 10% charcoal/dextran-treated serum for 48hrs prior to hormone treatment. Subsequently, cells were left untreated or treated with 10 nM aldosterone or 100 nM corticosterone for the indicated periods of time. Control cells were treated with ethanol at the same dilution used for treatments (1:1000).

### ChIP-seq and analysis

Cells were treated with vehicle, 10 nM aldosterone or 100 nM corticosterone for 1h. Chromatin crosslinking, preparation and immunoprecipitation was performed essentially as described (33). Briefly, chromatin crosslinking was performed using 1% formaldehyde added to culture medium for 5 min. After glycine quenching and washing with PBS, cells were recovered and chromatin extracted and sonicated (Bioruptor, Diagenode) to an average DNA length of 500 bp. For immunoprecipitation of GFP-MR, 600 μgs of chromatin were incubated with 25 µg anti-GFP antibody (Abcam #ab290).

The ChIP-Seq data were aligned to the mouse reference mm10 genome using Bowtie 2 with command Bowtie2 –p 8 –x bowtie2_ref/genome_prefix –U read1.fastq –S result.sam. Subsequent downstream analysis was performed using HOMER (70). Peaks in each dataset were called using the findPeaks function with style factor for TFs and the no treatment condition used as a control. Peak filtering was done with the following parameters; FDR<0.001, >5 FC over control, >5 FC over local background, and ntagThreshold >5 Peak clusters were identified by the mergePeaks command and sorted by cell type and treatment. Pre-defined motif searches were performed with findMotifsGenome.pl using -m known5.motif -mscore. Gene annotation of peaks used annotatePeaks.pl mm10 -gene.

### RNA Isolation, qPCR and RNA-seq analysis

Cells treated with vehicle, 10 nM aldosterone or 100 nM corticosterone for 2 h prior to RNA isolation. Total RNA was isolated using a commercially available kit (Macherey-Nagel NucleoSpin RNA isolation), which included an in-column DNase digestion step. Purified RNA was quantified using spectrophotometry and frozen in aliquots at -80°C. One aliquot was used to synthesize single-stranded cDNA starting from 1 µg of total RNA using a commercially available kit (iScript cDNA Synthesis Kit, Biorad).

RNA-seq included two to three biological replicates of each condition and used Illumina Novaseq with 150 bp stranded reads. RTA 2.4.11 was used for Base calling and Bcl2fastq 2.20 was used for demultiplexing allowing 1 mismatch. Cutadapt 1.18 was used for adapter removal and quality control. RNA-seq alignment to mouse mm10 genome was performed by STAR 2.70 using the default parameters with the following modifications: ‘--genomeDir mm10-125 --outSAMunmapped Within --outFilterType BySJout --outFilterMultimapNmax 20 --outFilterMismatchNmax 999 -- outFilterMismatchNoverLmax 0.04 --alignIntronMin 20 --alignIntronMax 1000000 -- alignMatesGapMax 1000000 --alignSJoverhangMin 8 --limitSjdbInsertNsj 2500000 -- alignSJDBoverhangMin 1 --sjdbScore 1 --sjdbFileChrStartEnd mm10-125/sjdbList.out.tab -- sjdbGTFfile UCSC_mm10_genes.gtf --peOverlapNbasesMin 10 --alignEndsProtrude 10 ConcordantPair’. All RNA-seq biological replicates correlated well with each other. Subsequent downstream analysis was performed using HOMER pipeline. Briefly, we obtained raw count data using analyzeRepeats.pl, and then the raw counts were normalized by default size factors from DESeq2 routine 23 provided via getDiffExpression.pl. We obtained differential genes using DESeq2, which fits negative binomial generalized linear models for each gene and uses the Wald test for significance testing, based on the criteria of a false discovery rate (FDR) cutoff <0.01 and absolute log2 fold change (FC) > 0.5 between no treatment and 2Hrs hormone treatment. We included only protein coding genes that are annotated in the RefSeq database and included no non-coding RNA species.

### Heatmap and aggregate plot generation

We used Deeptools to generate ChIP-seq and eRNA heatmaps and aggregate plots. We first generated read-normalized bigwig files from bam files using the bamCoverage -b [inputfile] -o [output.bigWig] -of bigwig --binSize 20 --effectiveGenomeSize 2652783500 --normalizeUsing RPGC. We generated matrix files using computeMatrix reference-point --referencePoint center -S [input.bigWig files] -R [peakfile.bed] -a 500 -o [matrix.gz] --sortRegions keep. We then generated heatmaps using plotHeatmap -m [matrix.gz] -o [HM.pdf] --sortRegions no --zMin --zMax -- refPointLabel "0 " --yAxisLabel "Tag Density". The eRNA heatmaps used merged replicate RNA bam files from the RNA seq data to make bigwig files. Bam files were merged using samtools.

### Single-molecule tracking

#### Transient transfections

GRKO or Parental cell lines were plated in complete medium in two-well LabTek II chamber slides. The next day, HaloTag MR or H2B constructs were transfected using jetOPTIMUS (Polyplus) following manufacturer’s protocol. After incubation for 4 hours with the jetOPTIMUS reaction mix, the medium was replaced with DMEM medium supplemented with charcoal/dextran-stripped FBS. 24 hours later, cells were incubated for 20 min with 5 nM of the cell-permeant HaloTag ligand Janelia Fluor 646 (JF_646_). After labeling, cells were washed three times for 15 min with phenol red-free DMEM media (Gibco) supplemented with charcoal/dextran-stripped FBS, followed by one last wash after 10 min, to remove unbound JF_646_. Cells were then treated with 10 nM Aldo or 100 nM Cort for 30 min before imaging.

#### Microscopy

All single-molecule tracking was performed on a custom-built HILO microscopy described previously (50). The microscope is equipped with a 150 X, 1.45 NA objective, (Olympus Scientific Solutions, Waltham, MA, USA), an Evolve 512 EM-CCD camera (Photometrics, Tucson, AZ, USA), a 647 nm laser (Coherent OBIS 647LX) and an Okolab stage-top incubator with 5% CO_2_ control. Images were collected every 200 ms with an exposure time of 10 ms and laser power of 0.85 mW at the objective. The pixel size for this microscope is 104 nm.

#### Tracking

Tracking was performed using TrackRecord v6, a custom MATLAB software freely available at Zenodo (https://doi.org/10.5281/zenodo.7558712) and has been described previously (71, 72). We allowed a maximum jump of 4 pixels, shortest track of 6 frames, and a gap of 1 frame. For details, see Wagh et al. (53).

#### Estimating the MSD distribution from single molecule trajectories

We calculate the self-part of the van Hove correlation function (vHc) *G_s_*(*r*, *τ*) = *A_s_*〈*δ*(*r_i_* −|*r*_*i*_(*t* + *τ*) – *r_i_*(*t*)|〉, from the single-molecule trajectories. Here ***r_i_*** is the position of the i^th^ nucleosome and *A_s_* = *∫ d*^2^*rG_s_*(*r*, *τ*) is a normalization constant. The vHc can be approximated as a superposition of Gaussian basis functions 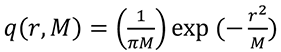 such that *G*_*s*_(*r*, *τ*) = *∫ P*(*M*, *τ*)*q*(*r*, *M*) *dM*. *P*(*M*) is the distribution of mean-squared displacements of the population of MR molecules. The Richardson-Lucy algorithm is then used the extract *P*(*M*) by iteratively fitting the vHc and updating the estimate of *P*(*M*). We refer the reader to references (51) and (53) for further details.

#### Classifying tracks into different mobility groups

Once the MSD distribution is calculated, the peaks in the distribution can be used to classify trajectories into different mobility groups. As shown in Figure S2, the local minima in the MSD distribution can be used to define four different mobility groups. The MSD of each track is then calculated at a timelag of 0.8 s, and by comparing this MSD to the four bins, the track is assigned to one of the mobility groups. The population fractions are calculated as the ratio of the number of tracks in a particular group to the total number of tracks.

### Data availability

The datasets produced in this study are available in the following databases:

- RNA-Seq data: Gene Expression Omnibus (GEO), accession number GSE232089.
- Chip-Seq data: Gene Expression Omnibus (GEO), accession number GSE232089.
- Single-molecule trajectories: Zenodo (https://doi.org/10.5281/zenodo.7846156).

## Acknowledgments

The authors thank the National Cancer Institute Advanced Technology Program Sequencing Facility for sequencing services. This research used the NIH high-performance computing systems (Biowulf) for genomics analyses. The researchers also thank Tatiana Karpova and David Ball of the Optical Microscopy Core at the NCI, NIH for assistance with the single molecule tracking experiments, and Diego M. Presman for his comments on the manuscript. Research was supported by grants from the Intramural Research Program of the National Institutes of Health, National Cancer Institute, Center for Cancer Research and by grant PID2019-105339RB-I00 (funded by MCIN/AEI/10.13039/501100011033 and “ERDF A way of making Europe”, MICINN, Spain). D.A.d.l.R. was partially supported by PRX18/00498 (funded by *Programa Estatal de Promoción del Talento y su Empleabilidad en I+D+i, Subprograma Estatal de Movilidad, del Plan Estatal de I+D+I,* MICINN, Spain). B.A.-P. was supported by pre-doctoral fellowship BES-2017- 082939 (funded by MCIN/AEI/ 10.13039/501100011033 and by “ESF Investing in your future”). A.U. was supported by NIH R35-145313 and NSF 2132922.

## Author Contributions

Conceptualization: TAJ, GF, GLH, DAdlR; Methodology: TAJ, GF, DAdlR; Formal analysis: TAJ, KW, AU; Investigation: TAJ, GF, BA-P, MK, DAdlR; Writing - original draft: TAJ, DAdlR; Writing - review & editing: TAJ, GF, KW, BA-P, AU, GLH, DAdlR; Visualization: TAJ, KW, DAdlR; Supervision: AU, GLH, DAdlR; Project administration: GLH, DAdlR; Funding acquisition: AU, GLH, DAdlR.

