## Supplemental Figures S1 and S2 for "The Glucocorticoid Receptor is Required for Efficient Aldosterone-Induced Transcription by the Mineralocorticoid Receptor"

**This PDF file includes:**

Figures S1 and S2

**Other supporting materials for this manuscript include the following:**

Tables S1 to S3

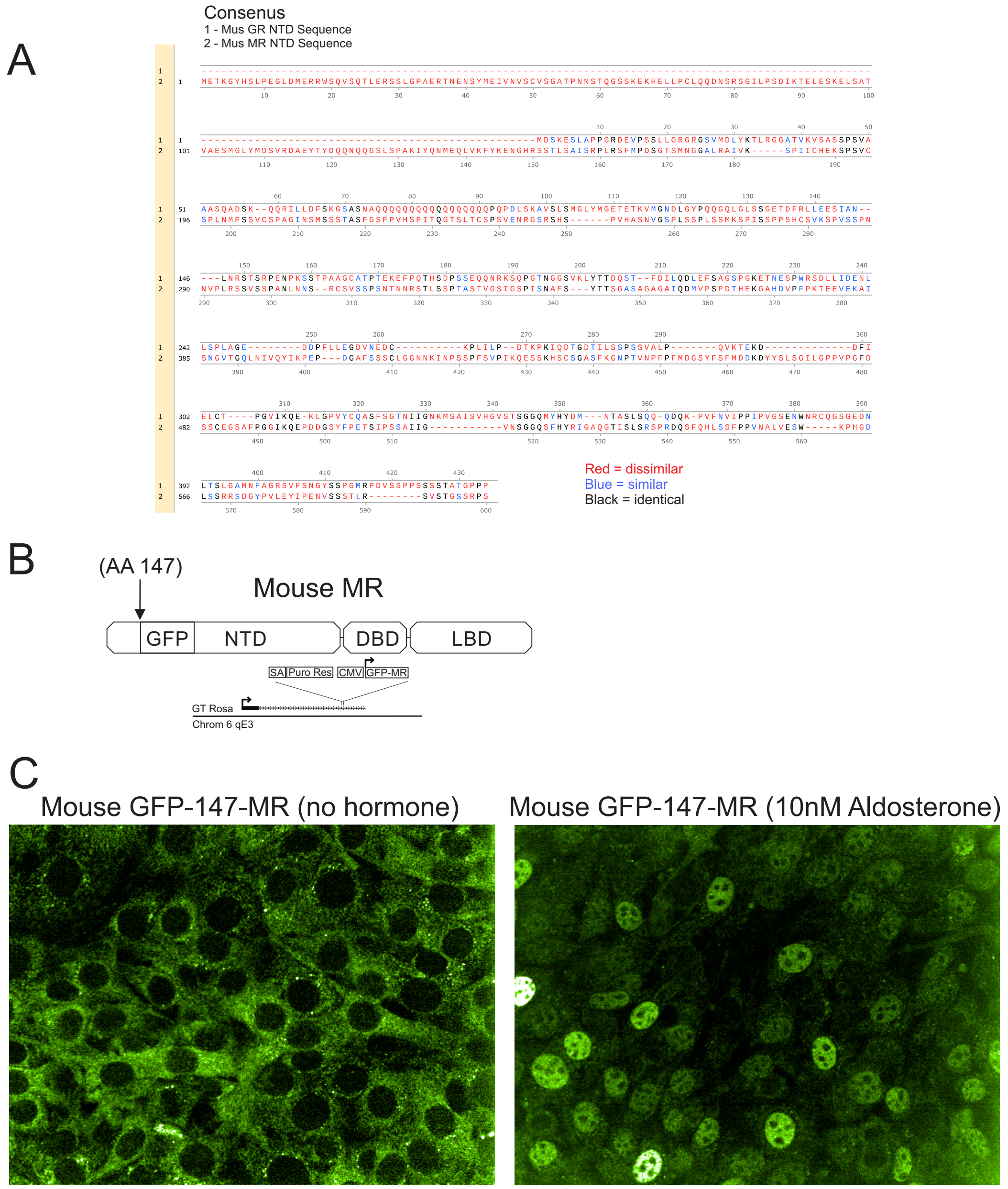


**Figure S1.** A. Sequence comparison between *M. musculus* MR and GR NTD. B. Schematic representation of M. musculus MR indicating the eGFP insertion site and the structure of the Donor-Rosa26_Puro_CMV-eGFP-MR vector. C. Representative confocal images showing eGFP-MR expression in a stable cell line and nuclear translocation after 1h 10 nM aldosterone treatment.

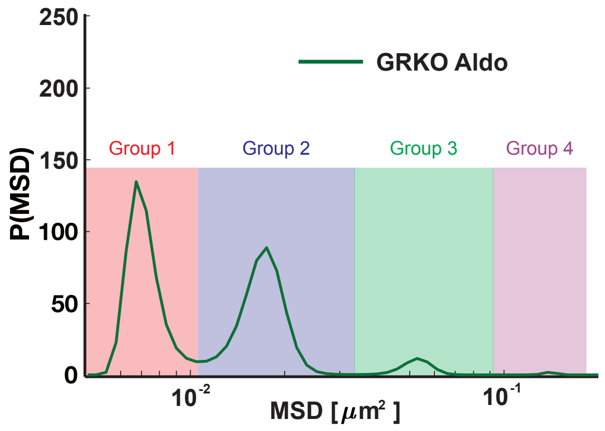


**Figure S2. Identification of different mobility groups from the MSD distribution.** A. representative mean squared displacement (MSD) distribution obtained by iteratively fitting the van Hove correlation function is shown. The local minima can be used to identify four distinct mobility groups.
